# Peripartum cardiomyopathy and hypertensive disorders of pregnancy and cardiovascular events among 1.6 million California pregnancies

**DOI:** 10.1101/404350

**Authors:** Rima Arnaout, Gregory Nah, Gregory M. Marcus, Zian H. Tseng, Elyse Foster, Ian Harris, Punag Divanji, Liviu Klein, Juan M. Gonzalez, Nisha I. Parikh

## Abstract

**Background:** Cardiovascular complications during and soon after pregnancy present an opportunity to assess risk for subsequent cardiovascular disease. We sought to determine whether peripartum cardiomyopathy and hypertensive disorder of pregnancy subtypes predict future myocardial infarction, heart failure, or stroke independent of one another and independent of other risks like gestational diabetes, preterm birth, and intrauterine growth restriction.

**Methods and Results:** The California Healthcare Cost and Utilization Project database was used to identify all hospitalized pregnancies from 2005-2009, with follow-up through 2011, for a retrospective cohort study. Pregnancies, exposures, covariates and outcomes were defined by ICD-9 codes. Among 1.6 million pregnancies (mean age 28y; median follow-up time to event 2.7y), 558 cases of peripartum cardiomyopathy, 123,603 cases of hypertensive disorders of pregnancy, 107,636 cases of gestational diabetes, 116,768 preterm births, and 23,504 cases of intrauterine growth restriction were observed. Using multivariable Cox proportional hazards models, peripartum cardiomyopathy was independently associated with a 13.0-fold increase in myocardial infarction [95%CI, 4.1-40.9], a 39.2-fold increase in heart failure [95%CI, 30.0-51.9], and a 7.7-fold increase in stroke [95%CI, 2.4-24.0]. Hypertensive disorders of pregnancy were associated with a 1.4 [95%CI, 1.0-2.0] to 7.6 [95%CI, 5.4-10.7] fold higher risk of myocardial infarction, heart failure, and stroke. Gestational diabetes, preterm birth, and intrauterine growth restriction had more modest associations with CVD.

**Conclusions:** These findings support close monitoring of women with cardiovascular pregnancy complications for prevention of early subsequent cardiovascular events and further study of mechanisms underlying their development.

## Introduction

Cardiovascular disease (CVD) is a significant and often underappreciated cause of morbidity and mortality in women(1). Pregnancy can often “unmask” CVD in women(2,3). In the estimated 85 percent of women who experience a pregnancy during their lifetime, cardiovascular complications during pregnancy and childbirth therefore present an opportunity to assess risk for later CVD(4). Indeed, leveraging early phenotypes to identify those at risk for CVD as soon as possible is an aspirational goal for both precision medicine(5,6) and for primary CVD disease prevention(7,8).

Peripartum cardiomyopathy (PPCM) and hypertensive disorders of pregnancy (HDP) are two major cardiovascular complications of pregnancy and have been reported to share a common underlying pathophysiology in animal studies(9-11). PPCM is characterized by the sudden onset of maternal heart failure presenting either in the last month of pregnancy or in the first five months postpartum. Hypertension in pregnancy is defined as a blood pressure equal to or greater than 140/90 mmHg; gestational onset is defined at or after 20 weeks’ gestation (**Online Table 2**).

We hypothesized that specific subtypes of PPCM and HDP may carry different CVD risks(12),(13). Recent data from a Danish nationwide register–based cohort demonstrated that specific subtypes of HDP (i.e. severe preeclampsia, moderate preeclampsia and gestational hypertension) conferred differing risks of incident cardiomyopathy(14). Still, prior studies have not fully elucidated the independent risks of PPCM and HDP subtypes on specific CVD outcomes [i.e. myocardial infarction (MI), heart failure (HF), and stroke]. The availability of a large cohort of women directly representative of the general population of California in the California Healthcare Cost and Utilization Project (HCUP) allowed us to determine the risks of PPCM and HDP subtypes along a spectrum of chronicity and severity for subsequent MI, HF, and stroke, while independently adjusting for other demographic and pregnancy-related factors that are known to be associated with later maternal CVD [i.e. gestational diabetes mellitus (GDM), preterm delivery and intrauterine growth restriction (IUGR)](15-17).

## Methods

### Study Sample

We used the California Healthcare Cost and Utilization Project (HCUP) database (https://www.hcup-us.ahrq.gov/), which provides state-specific data for all inpatient, emergency, and ambulatory visits. HCUP contains information collected as part of medical billing, including patient demographics, International Classification of Diseases, Ninth Revision (ICD-9) diagnoses, expected payer, dates of admission and discharge, and follow-up. We identified the first delivery from all patients ages 18 and older in HCUP from 2005-2009 by ICD-9 code (n=1,686,601). During the 2005-2009 study period, after excluding those with a non-California residence (n=2,734), missing information on age (n=2,742) or race/ethnicity (n=11,434), data were available in California HCUP for 1,669,691 pregnancies. Among those 1,669,691 remaining pregnancies we excluded those with pre-existing MI, HF, stroke, and congenital(18) or valvular heart disease using ICD-9 codes (n=7,646), leaving 1,662,045 eligible pregnancies for this study. Congenital heart disease diagnoses were classified according to methods previously described.(18) The number of pregnancies in our study cohort represents 7.8% of all pregnancies in the United States within the same time period(19). Analyses were performed in accordance with the HCUP Data Use Agreement. To preserve patient anonymity, any groups in which there were fewer than 10 patients are listed in Tables as <10.

### Exposures

After pregnancies were identified, we defined instances of PPCM, HDP subtypes, GDM, preterm birth, and IUGR by ICD-9 code (**Online Table 1**). We considered both principal and secondary diagnoses to identify these pregnancy exposures.

**Table 1.**
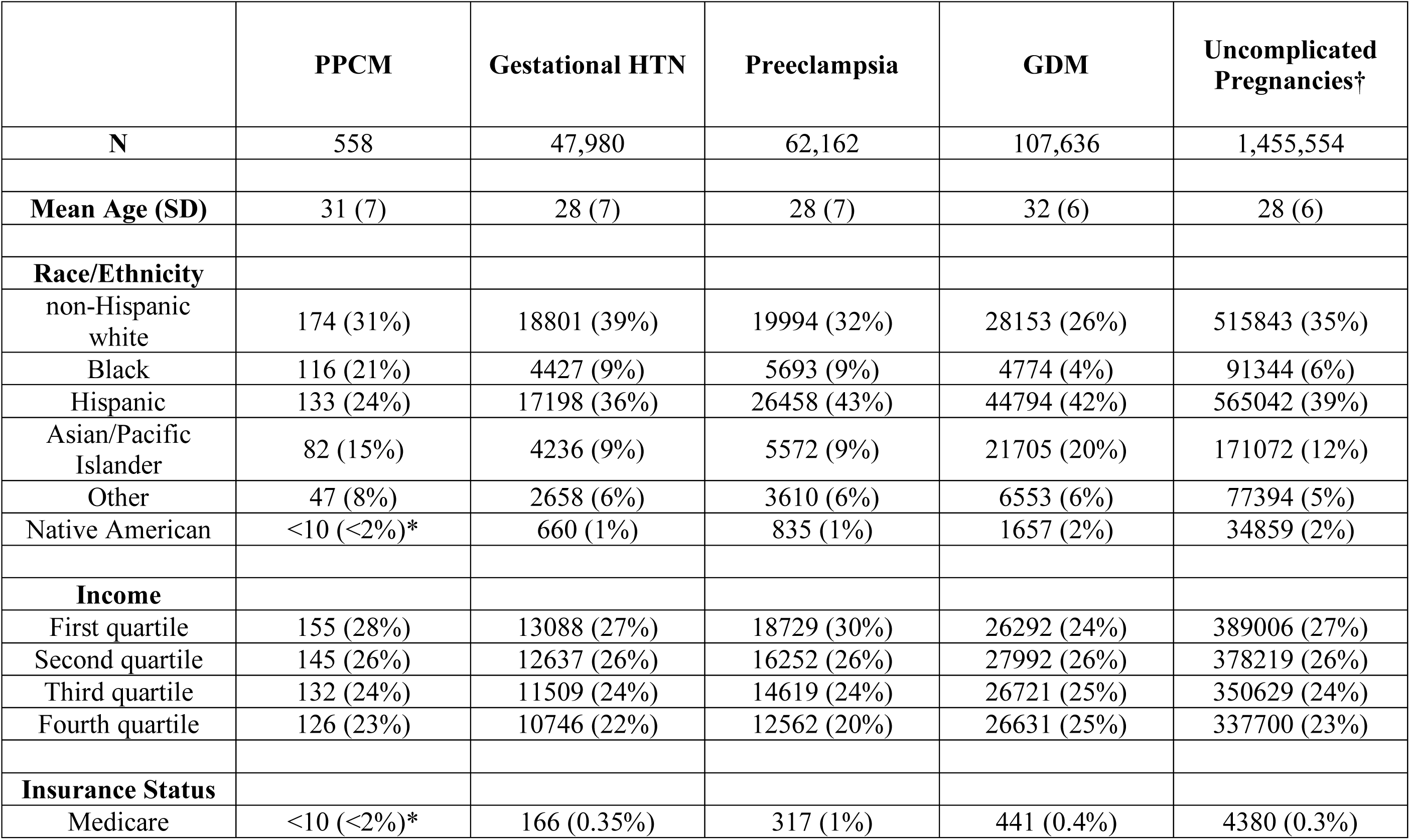
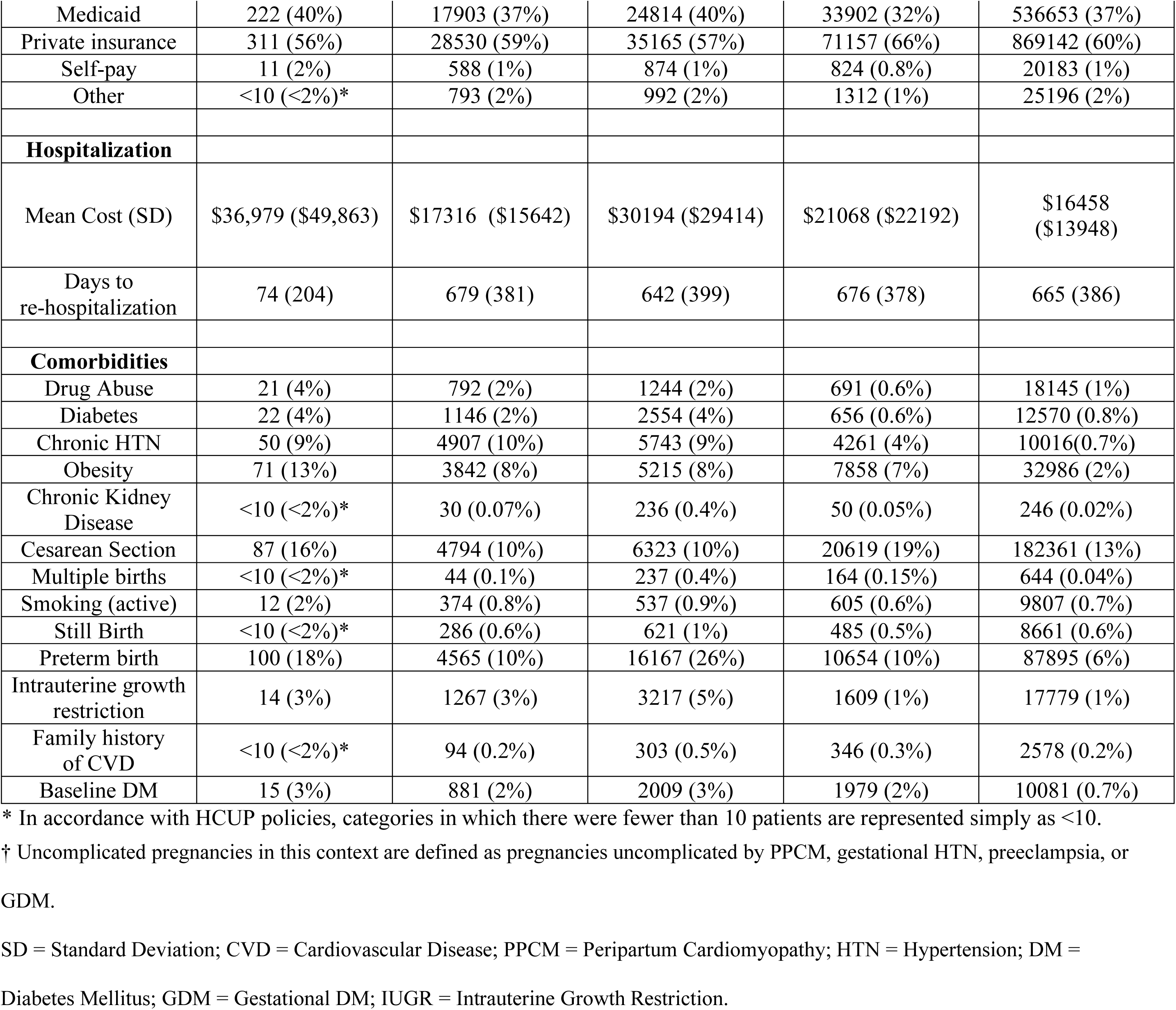
Patient characteristics. Baseline characteristics for patients included in the analysis; those with preexisting MI, heart failure, stroke, and congenital or valvular heart disease were excluded.

|  | **PPCM** | **Gestational HTN** | **Preeclampsia** | **GDM** | **Uncomplicated Pregnancies†** |
| --- | --- | --- | --- | --- | --- |
| **N** | 558 | 47,980 | 62,162 | 107,636 | 1,455,554 |
| **Mean Age (SD)** | 31 (7) | 28 (7) | 28 (7) | 32 (6) | 28 (6) |
| **Race/Ethnicity** |  |  |  |  |  |
| non-Hispanic white | 174 (31%) | 18801 (39%) | 19994 (32%) | 28153 (26%) | 515843 (35%) |
| Black | 116 (21%) | 4427 (9%) | 5693 (9%) | 4774 (4%) | 91344 (6%) |
| Hispanic | 133 (24%) | 17198 (36%) | 26458 (43%) | 44794 (42%) | 565042 (39%) |
| Asian/Pacific Islander | 82 (15%) | 4236 (9%) | 5572 (9%) | 21705 (20%) | 171072 (12%) |
| Other | 47 (8%) | 2658 (6%) | 3610 (6%) | 6553 (6%) | 77394 (5%) |
| Native American | <10 (<2%)* | 660 (1%) | 835 (1%) | 1657 (2%) | 34859 (2%) |
| **Income** |  |  |  |  |  |
| First quartile | 155 (28%) | 13088 (27%) | 18729 (30%) | 26292 (24%) | 389006 (27%) |
| Second quartile | 145 (26%) | 12637 (26%) | 16252 (26%) | 27992 (26%) | 378219 (26%) |
| Third quartile | 132 (24%) | 11509 (24%) | 14619 (24%) | 26721 (25%) | 350629 (24%) |
| Fourth quartile | 126 (23%) | 10746 (22%) | 12562 (20%) | 26631 (25%) | 337700 (23%) |
| **Insurance Status** |  |  |  |  |  |
| Medicare | <10 (<2%)* | 166 (0.35%) | 317 (1%) | 441 (0.4%) | 4380 (0.3%) |
| Medicaid | 222 (40%) | 17903 (37%) | 24814 (40%) | 33902 (32%) | 536653 (37%) |
| Private insurance | 311 (56%) | 28530 (59%) | 35165 (57%) | 71157 (66%) | 869142 (60%) |
| Self-pay | 11 (2%) | 588 (1%) | 874 (1%) | 824 (0.8%) | 20183 (1%) |
| Other | <10 (<2%)* | 793 (2%) | 992 (2%) | 1312 (1%) | 25196 (2%) |
| **Hospitalization** |  |  |  |  |  |
| Mean Cost (SD) | \$36,979 (\$49,863) | \$17316 (\$15642) | \$30194 (\$29414) | \$21068 (\$22192) | \$16458 (\$13948) |
| Days to re-hospitalization | 74 (204) | 679 (381) | 642 (399) | 676 (378) | 665 (386) |
| **Comorbidities** |  |  |  |  |  |
| Drug Abuse | 21 (4%) | 792 (2%) | 1244 (2%) | 691 (0.6%) | 18145 (1%) |
| Diabetes | 22 (4%) | 1146 (2%) | 2554 (4%) | 656 (0.6%) | 12570 (0.8%) |
| Chronic HTN | 50 (9%) | 4907 (10%) | 5743 (9%) | 4261 (4%) | 10016(0.7%) |
| Obesity | 71 (13%) | 3842 (8%) | 5215 (8%) | 7858 (7%) | 32986 (2%) |
| Chronic Kidney Disease | <10 (<2%)* | 30 (0.07%) | 236 (0.4%) | 50 (0.05%) | 246 (0.02%) |
| Cesarean Section | 87 (16%) | 4794 (10%) | 6323 (10%) | 20619 (19%) | 182361 (13%) |
| Multiple births | <10 (<2%)* | 44 (0.1%) | 237 (0.4%) | 164 (0.15%) | 644 (0.04%) |
| Smoking (active) | 12 (2%) | 374 (0.8%) | 537 (0.9%) | 605 (0.6%) | 9807 (0.7%) |
| Still Birth | <10 (<2%)* | 286 (0.6%) | 621 (1%) | 485 (0.5%) | 8661 (0.6%) |
| Preterm birth | 100 (18%) | 4565 (10%) | 16167 (26%) | 10654 (10%) | 87895 (6%) |
| Intrauterine growth restriction | 14 (3%) | 1267 (3%) | 3217 (5%) | 1609 (1%) | 17779 (1%) |
| Family history of CVD | <10 (<2%)* | 94 (0.2%) | 303 (0.5%) | 346 (0.3%) | 2578 (0.2%) |
| Baseline DM | 15 (3%) | 881 (2%) | 2009 (3%) | 1979 (2%) | 10081 (0.7%) |

\* In accordance with HCUP policies, categories in which there were fewer than 10 patients are represented simply as <10.
† Uncomplicated pregnancies in this context are defined as pregnancies uncomplicated by PPCM, gestational HTN, preeclampsia, or GDM. SD = Standard Deviation; CVD = Cardiovascular Disease; PPCM = Peripartum Cardiomyopathy; HTN = Hypertension; DM = Diabetes Mellitus; GDM = Gestational DM; IUGR = Intrauterine Growth Restriction.

We further divided PPCM into diagnoses made pre-partum (PPCM diagnoses already present at delivery) and those made post-partum (PPCM diagnosis appearing within 5 months post-delivery). We defined HDP as one of the following: chronic HTN, gestational HTN, preeclampsia, chronic HTN with gestational HTN, chronic HTN with preeclampsia and no HTN or PPCM (referent) (**Online Table 2**).

**Table 2.** Multivariable-adjusted associations between pregnancy complications and myocardial infarction, heart failure, and stroke.

| All eligible pregnancies<br>N = 1,662,045 | Myocardial<br>Infarction<br>MV adjusted HR<br>(95% CI)<br>n = 558 | Heart Failure<br>MV adjusted HR<br>(95% CI)<br>n = 3,284 | Stroke<br>MV adjusted HR<br>(95% CI)<br>n = 973 |
| --- | --- | --- | --- |
| PPCM<br>n = 558<br>1.66 per 1000 person-years | 13.0 (4.1-40.9)<br>n = <10*<br>N/A | 39.2 (30.0 – 51.9)<br>n = 56<br>2.19 per 1000<br>person-years | 7.7 (2.4-24.0)<br>n = <10*<br>N/A |
| No PPCM (referent)<br>n=1,661,487<br>0.58 per 1000 person-years | 1.0<br>n=549<br>0.98 per 1000<br>person-years | 1.0<br>n=3228<br>1.22 per 1000<br>person-years | 1.0<br>n=964<br>1.01 per 1000<br>person-years |
| HP subtypes: |  |  |  |
| Chronic HTN<br>n = 12,458<br>0.54 per 1000 person-years | 6.3 (4.6-8.7)<br>n = 47<br>0.90 per 1000<br>person-years | 3.9 (3.3-4.7)<br>n = 152<br>1.12 per 1000<br>person-years | 3.4 (2.4-4.8)<br>n = 37<br>0.88 per 1000<br>person-years |
| *De novo*<br>gestational HTN<br>n = 43,073<br>5.98 per 1000 person-years | 2.3 (1.6-3.4)<br>n = 28<br>0.98 per 1000<br>person-years | 2.5 (2.1-2.9)<br>n = 188<br>1.39 per 1000<br>person-years | 1.4 (1.0-2.0)<br>n = 33<br>1.15 per 1000<br>person-years |
| *De novo*<br>preeclampsia<br>n = 56,419 | 2.5 (1.9-3.4)<br>n = 49 | 3.0 (2.7-3.4)<br>n = 355 | 2.3 (1.8-3.0)<br>n = 79 |
| 0.59 per 1000 person-years | 0.89 per 1000 person-years | 1.69 per 1000 person-years | 1.08 per 1000 person-years |
| Chronic HTN & Gestational HTN<br>n = 4,907<br>0.54 per 1000 person-years | 2.0 (0.8-4.8)<br>n = <10*<br>N/A | 3.3 (2.5-4.5)<br>n = 45<br>1.16 per 1000 person-years | 3.0 (1.7-5.4)<br>n = 12<br>1.67 per 1000 person-years |
| Chronic HTN & Preeclampsia<br>n = 5,743<br>0.61 per person-years | 5.0 (3.1-8.0)<br>n = 22<br>1.01 per 1000 person-years | 5.4 (4.5-6.6)<br>n = 135<br>1.36 per 1000 person-years | 7.6 (5.4-10.7)<br>n = 43<br>1.03 per 1000 person-years |
| No HTN/preeclampsia/gestational htn (referent)<br>n = 1,539,445<br>0.57 per 1000 person-years | 1.0<br>n=407<br>1.00 per 1000 person-years | 1.0<br>n = 2,409<br>1.17 per 1000 person-years | 1.0<br>n = 769<br>1.01 per 1000 person-years |
| Gestational DM<br>n = 107,636<br>0.59 per 1000 person-years | 1.0 (0.8-1.4)<br>n=57<br>0.88 per 1000 person-years | 1.3 (1.1-1.4)<br>n = 320<br>1.36 per 1000 person-years | 1.42(1.2-1.8)<br>n = 103<br>1.09 per 1000 person-years |
| No Gestational DM (referent)<br>n=1,554,409<br>0.58 per 1000 person-years | 1.0<br>n=501<br>1.00 per 1000 person-years | 1.0<br>n=2964<br>1.22 per 1000 person-years | 1.0<br>n=870<br>1.01 per 1000 person-years |
| IUGR<br>n = 23,504<br>0.59 per 1000 person-years | 1.6 (1.1-2.5)<br>n = 23<br>1.11 per 1000 person-years | 1.1 (0.9-1.4)<br>n = 86<br>1.22 per 1000 person-years | 1.5 (1.0-2.1)<br>n = 30<br>1.09 per 1000 person-years |
| No IUGR (referent)<br>n=1,638,541 | 1.0<br>n=535 | 1.0<br>n=3198 | 1.0<br>n=943 |
| 0.58 per 1000 person-years | 0.98 per 1000 person-years | 1.23 per 1000 person-years | 1.02 per 1000 person-years |
| Preterm birth<br>n=116,768<br>0.58 per 1000 person-years | 1.8 (1.4-2.3)<br>n=105<br>1.05 per 1000 person-years | 1.6 (1.4-1.7)<br>n=532<br>1.25 per 1000 person-years | 1.4 (1.1-1.6)<br>n=134<br>1.05 per 1000 person-years |
| Preterm birth(referent)<br>n=1,545,277<br>0.58 per 1000 person-years | 1.0<br>n=453<br>0.97 per 1000 person-years | 1.0<br>n=2752<br>1.23 per 1000 person-years | 1.0<br>n=839<br>1.01 per 1000 person-years |

\* In accordance with HCUP policies, categories in which there were fewer than 10 patients are represented simply as <10.
MV = Multivariate; HR = Hazard Ratio; CI = Confidence Interval; HDP = Hypertensive Disorders of Pregnancy; PPCM = Peripartum
Cardiomyopathy; HTN = Hypertension; DM = Diabetes Mellitus; IUGR = Intrauterine Growth Restriction.

### Outcomes

Primary CVD outcomes were admissions for MI, HF, and stroke defined by ICD-9 codes (**Online Table 1**). MI subtypes were defined as MI with CAD (MICAD) and MI with non-obstructive coronary arteries (MINOCA). MINOCA includes MI from stress cardiomyopathy, hypercoagulable state, coronary artery dissection, coronary artery anomaly, and coronary vasospasm.(20) MINOCA was defined using its ICD-9 code as well as 1) having had a myocardial infarction using ICD-9 codes 410.00-410.92; 2) having had a cardiac catheterization or computed tomography coronary angiogram (inpatient or outpatient, defined by CPT code) within 7 days of index event; and 3) having had an ICD-9 code for a MINOCA subtype. ICD-9 codes used for MINOCA subtypes were 413.1 for coronary vasospasm; 414.12 for coronary artery dissection; 429.83 for Takotsubo/stress cardiomyopathy; 289.81 for hypercoagulable state; and 746.85 for coronary anomaly.

Stroke was further subdivided into ischemic, embolic, and hemorrhagic etiologies. Among our study cohort, systolic vs diastolic HF, and left vs right HF were not well specified and thus we were unable to distinguish among these HF subtypes. ICD-9 codes defining the outcomes are found in **Online Table 1**.

### Covariates

We adjusted for covariates known to be associated with PPCM, hypertension, CVD, and peripartum morbidity and mortality(21) which could potentially confound the associations we were interested in studying. Covariates included age, race, insurance status, median household income, chronic kidney disease, pre-existing diabetes, obesity, drug abuse, smoking, multiple gestations, as well as each of the exposures. Patients with preexisting (prior to pregnancy) HTN were considered to have baseline chronic hypertension and were included within the HDP variable; therefore, we did not include HTN as a covariate.

### Statistical Analyses

*Primary analyses.* We considered our baseline period in which initial delivery occurred from 2005-2009. Follow-up began 6 months after delivery in order to avoid double-counting an exposure and outcome, and it lasted until the end of 2011. Descriptive statistics were performed including n’s, means and percentages with standard deviations and interquartile ranges as appropriate among all participants and within PPCM and HDP subtypes. We tested the assumption of proportionality of hazards for PPCM and HDP subtypes and MI, HF, and stroke. We found that one of the five HDP subtypes, *de novo* preeclampsia, violated the assumption (p=0.02). Therefore, in the case of *de novo* preeclampsia, our hazard ratio estimates represent an average across the length of study follow-up. We employed multivariable Cox proportional hazards models to determine the association between PPCM and HDP and subsequent CVD, adjusting for the potential confounders listed in the covariates section above. We considered GDM, preterm delivery and IUGR as secondary exposures and considered these in multivariable models adjusting for PPCM and HDP.

### Secondary analyses

It is known that certain pregnancy complications beget further comorbidities: for example, hypertensive disorders of pregnancy can be risk factors for PPCM(14). *Overlap.* We assessed pregnancy exposures for overlap. Similarly, we assessed for overlap among our CVD outcomes of interest. Finally, since specific subtypes of MI and stroke may be more common in younger individuals and among women, we analyzed outcomes by subtype. For example, we assessed how many of the MI’s in our population were due to MICAD vs MINOCA, and how many of the incident strokes were from hemorrhagic versus ischemic versus embolic etiologies. As mentioned above, ICD-9 codes for HF subtypes were not specified in the study cohort. We also considered PPCM as intra-or postpartum in order to determine whether timing of PPCM affected CVD outcomes. *Stratification*. In addition to our original multivariable Cox proportional hazards model (see Primary analyses), we re-ran the model i) stratifying on quintiles of age rather than using age as a covariate and ii) stratifying by preterm birth and PPCM, preeclampsia and PPCM, and IUGR and PPCM.

All analyses were performed using SAS version 9.4X. A p-value of 0.05 or less was considered statistically significant. Certification to use de-identified HCUP data was obtained from the University of California, San Francisco Committee on Human Research. No informed consent was required.

## Results

### Characteristics of study population

Of the 1,662,045 study participants who delivered from 2005-2009 in HCUP-CA, 579,656 (35%) were non-Hispanic white women, 105,489 (6%) were African-American, 648,602 (39%) were Hispanic, and 200,908 (12%) were Asian or Pacific Islanders. 20,765 (1.2%) of pregnancies were complicated by drug abuse, and 11,248 (0.7%) of pregnancies were complicated by active smoking. 330, 946 deliveries were second pregnancies within the study period. In terms of additional risk factors for cardiovascular complications, 23,108 (1.4%) of pregnant women had pre-existing hypertension, 16,806 (1.0%) had pre-existing diabetes, and 538 (0.3%) had chronic kidney disease. Notably, the prevalence of African Americans, chronic HTN, and obesity was higher in the PPCM group than in the general population. The prevalence of Asian and Pacific Islanders was higher in the GDM group. **Table 1** summarizes these and other characteristics of the study cohort.

### Prevalence of pregnancy exposures in study population

There were 558 cases of PPCM, 97 of which were diagnosed before delivery, and 461 of which were diagnosed postpartum. There were 49,114 cases of gestational HTN and 62,162 cases of preeclampsia; broken down by HDP subtypes of interest, there were 12,458 cases of isolated chronic HTN, 43,073 cases of *de novo* gestational HTN, 56,419 cases of *de novo* preeclampsia, 4,907 pregnancies with both chronic HTN and superimposed gestational HTN, and 5,743 pregnancies with chronic HTN and superimposed preeclampsia (**Table 2**). There were 107,636 cases of GDM, 23,504 cases of IUGR, and 116,768 cases of preterm birth. Median follow-up time to event was 2.68 years, ranging from 1 day to 6.84 years, resulting in 0.99, 1.23, and 1.02 cases of MI, HF, and stroke, respectively, per 1000 person-years. **Online Table 3** demonstrates the overlap and correlations among pregnancy exposure categories. Preterm birth and preeclampsia were the most highly correlated (Kendall-Tau correlation coefficient 0.146) whereas IUGR and peripartum cardiomyopathy had the lowest correlation (Kendall-Tau correlation coefficient 0.0017) among pregnancy exposures studied.

**Table 3.** Number of patients in study cohort with CVD outcomes of interest by outcome subtype.

|  | MI |  | Stroke |  |  | Heart Failure |
| --- | --- | --- | --- | --- | --- | --- |
|  | MICAD | MINOCA | Embolic | Ischemic | Hemorrhagic |  |
| PPCM (n=558) | <10* | <10* | <10* | <10* | 0 | 56 |
| Chronic HTN (n=12,458) | 14 | <10* | <10* | 28 | <10* | 152 |
| *De Novo* Gestational HTN (n=43,073) | <10* | <10* | <10* | 22 | <10* | 188 |
| *De Novo* Preeclampsia (n=56,419) | 16 | <10 | <10 | 60 | 14 | 355 |
| Chronic HTN & Gestational HTN (n=4,907) | <10* | 0 | <10* | <10* | <10* | 45 |
| Chronic HTN & Preeclampsia (n=5,743) | <10* | 0 | <10* | 35 | <10* | 135 |
| Gestational DM (n=107,636) | 24 | <10* | 12 | 62 | 29 | 320 |
| Preterm Birth (n=116,768) | 42 | 15 | 13 | 96 | 25 | 532 |
| IUGR (n=23,504) | <10* | <10* | <10* | 17 | <10* | 86 |
| Total | 126 | 40 | 53 | 333 | 91 | 1251 |

\* In accordance with HCUP policies, categories in which there were fewer than 10 patients are represented simply as <10.
MI = Myocardial Infarction; MICAD = MI with obstructive coronary artery disease; MINOCA = MI without obstructive coronary artery disease; PPCM = Peripartum Cardiomyopathy; HTN = Hypertension; DM = Diabetes Mellitus; IUGR = Intrauterine Growth Restriction.

### Multivariable models demonstrated associations between exposures and future cardiovascular risk

Among all women hospitalized during pregnancy from 2005-2009, there were 558 subsequent MIs, 3284 subsequent cases of HF, and 973 subsequent strokes. **Table 2** shows stroke, MI, and HF event numbers by pregnancy exposure category. Of the 330,946 pregnancies (of the total ~1.6 million pregnancies) which were second pregnancies within the study period, 24 had MI (24 of 558), 133 had HF (133 of 3284), and 37 had stroke (37 of 973). Using multivariable Cox proportional hazards models adjusting for all exposures and covariates, we found that PPCM was independently associated with a 13.0-fold increased risk of MI (95% CI 4.1-40.9), a 39.2-fold increased risk of HF (95% CI 30.0 – 51.9), and a 7.7-fold increased risk of stroke (95% CI 2.4-24.0; **Figure 1; Table 2).** The hazard ratios were similar for PPCM diagnosed at or before delivery when compared to PPCM diagnosed in the first five months postpartum **(Figure 1)**.

**Figure 1.**
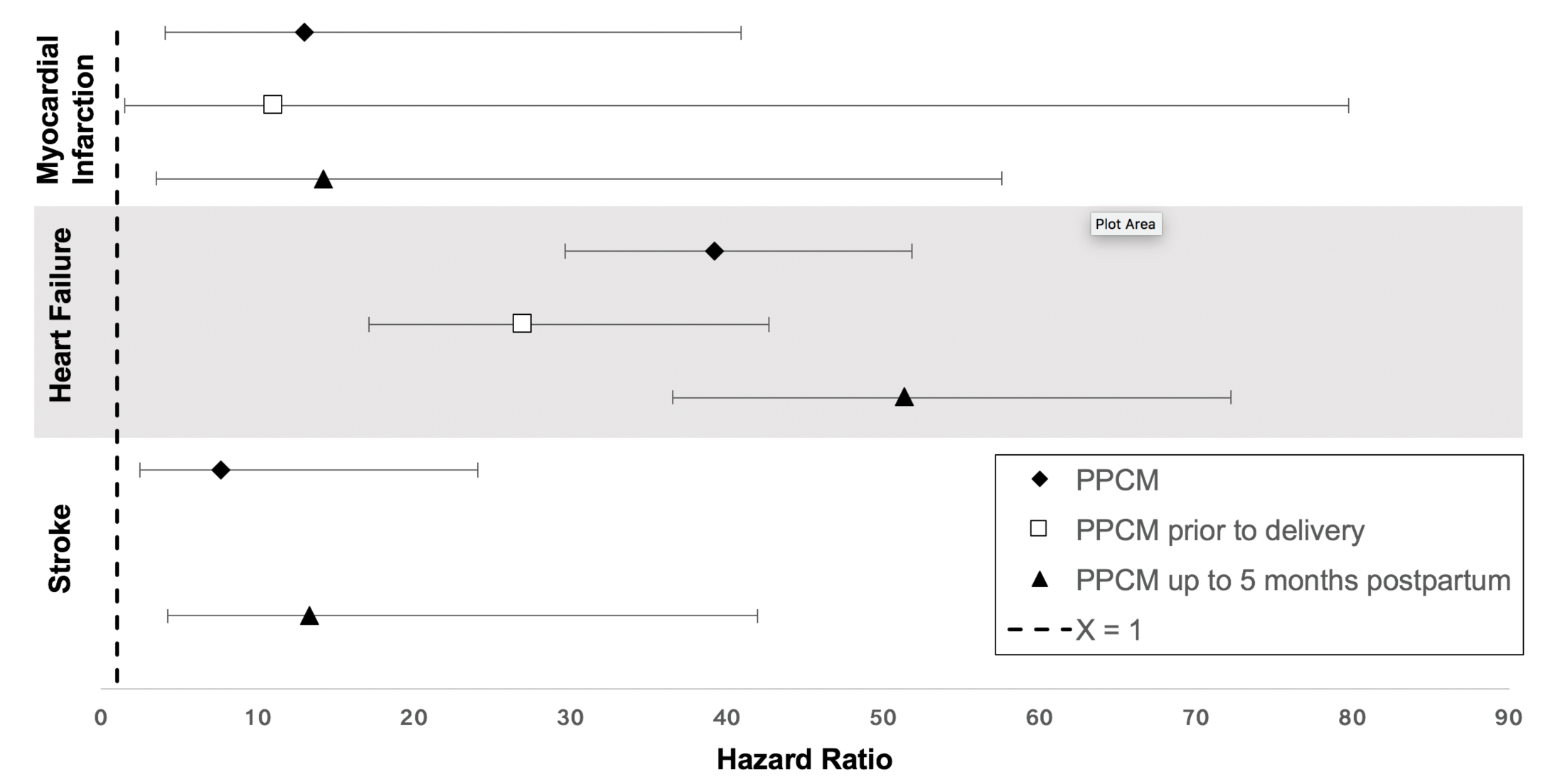
Adjusted associations between peripartum cardiomyopathy and myocardial infarction, heart failure, and stroke. Hazard ratios for all PPCM (black diamond), as well as PPCM diagnosed prior to delivery (open square) and postpartum (black triangle). Covariates included age, race, insurance status, median household income, chronic kidney disease, preexisting diabetes, obesity, drug abuse, smoking, multiple gestations, as well as for each of the exposures (PPCM, HDP subtypes, GDM, IUGR, and preterm birth).

We investigated cardiovascular risks according to hypertensive disorder of pregnancy subtypes of differing chronicity and severity. We examined five subtypes: chronic hypertension alone, *de novo* gestational hypertension, *de novo* preeclampsia, chronic hypertension with superimposed gestational hypertension, and chronic hypertension with superimposed preeclampsia **(Online Table 2)**. These subtypes carried a 2.3-6.3-fold increased risk for MI, a 2.5-5.4-fold increased risk for HF, and a 1.4-7.6-fold increased risk for stroke **(Figure 2; Table 2).** Although there was some overlap in 95% confidence intervals, chronic HTN alone, and with superimposed preeclampsia, carried the highest risks of MI, HF and stroke compared with other HDP subtypes. Chronic HTN with superimposed gestational hypertension did not demonstrate a statistically significant increased risk for MI; the n in this group was less than 10.

**Figure 2.**
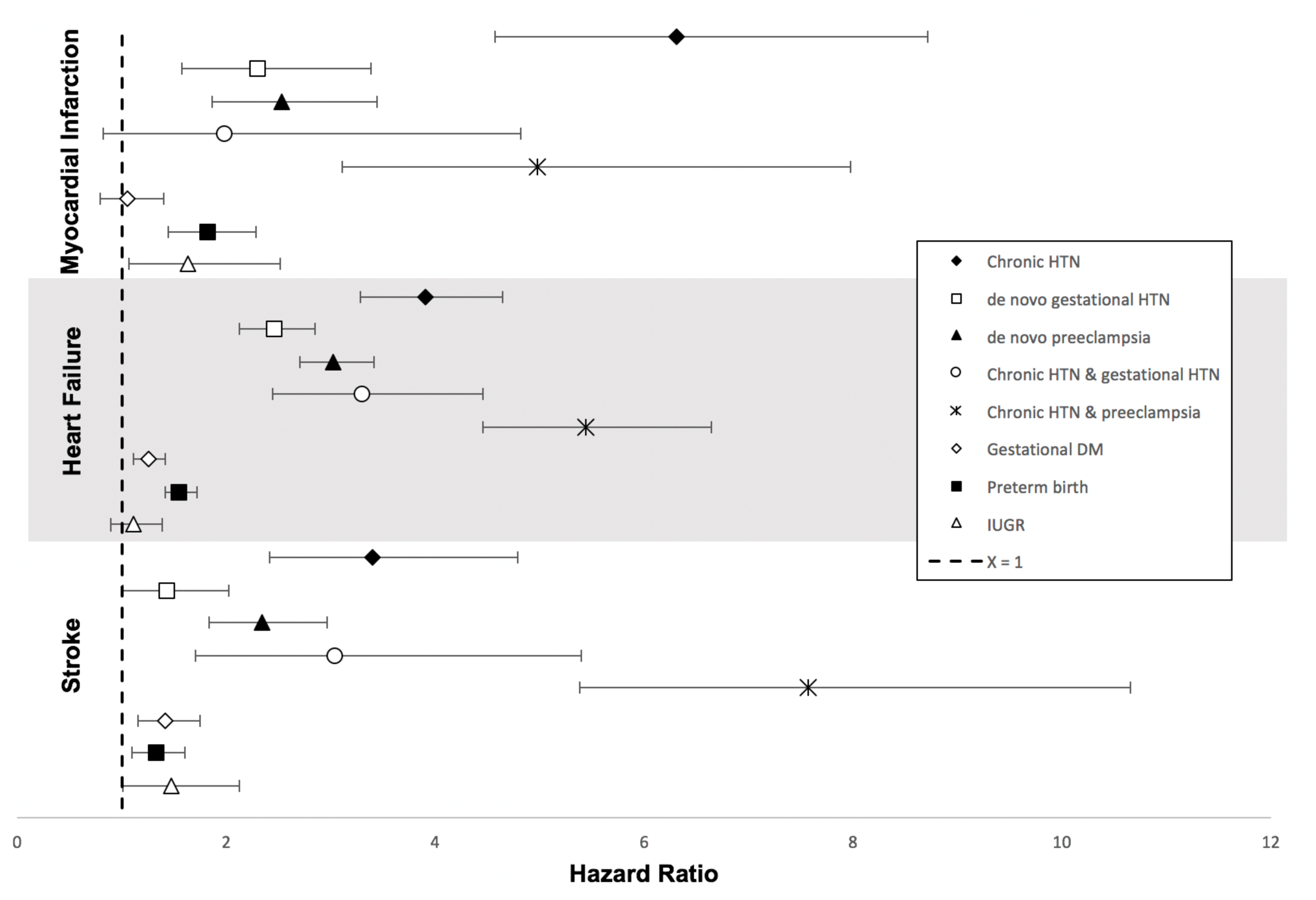
Adjusted associations between hypertensive disorder of pregnancy subtypes and myocardial infarction, heart failure, and stroke. Hazard ratios for HDP subtypes, GDM, IUGR and preterm birth and MI, HF, and stroke. Covariates included age, race, insurance status, median household income, chronic kidney disease, pre-existing diabetes, obesity, drug abuse, smoking, multiple gestations, as well as for each of the exposures (PPCM, HDP subtypes, GDM, IUGR, and preterm birth).

All of our models included additionally included GDM, preterm birth, and IUGR. IUGR was related to risk of MI but not HF or stroke [HR for MI=1.6 (95% CI: 1.0-2.5)]. Preterm delivery was significantly associated with MI, stroke and HF (Figure 2; Table 2). GDM was associated with risks of stroke and HF but not with MI **(Figure 2; Table 2).** Overall, these three pregnancy exposures conferred comparatively lower risks for cardiovascular diseases as compared to PPCM and HDP.

### Tests for residual confounding in the multivariable model show robust associations between exposures and outcomes

Age is a well-known and strong predictor both for both cardiovascular complications of pregnancy and for our outcomes of interest; (21,22) and increased prevalence of exposures and outcomes by age was again demonstrated in our study (**Online Table 4**). We therefore sought to test for residual confounding by age in our multivariable model. We compared hazard ratios derived from our original model to those obtained using an alternate approach stratifying on quintiles of age rather than using age as a covariate (thereby avoiding the assumptions of the proportional hazards model). We found that this alternate approach did not change hazard ratio point estimates or 95% confidence intervals by more than a point (data not shown).

We also ran additional models stratifying by preterm birth and PPCM, preeclampsia and PPCM, and IUGR and PPCM, and did not find substantial differences in hazard ratio point estimates or 95% confidence intervals (data not shown).

### Secondary analyses

With respect to exposures, excepting the HDP subtypes of gestational HTN & chronic HTN and preeclampsia & chronic HTN, where clinical overlap was present by design (10.3% and 9.24% overlap, respectively), there was notable overlap between PPCM and preeclampsia (26.1%), preeclampsia and preterm birth (26.0%) between chronic HTN and preterm birth (18.8%). Overlap among all other categories ranged from 1.49% to 10.6% (**Online Table 3**).

With respect to outcomes, PPCM is known to be a risk factor for HF but has not previously been described as a risk factor for MI or stroke (**Figure 1, Table 2**). To determine whether there was overlap in MI, HF, and stroke outcome diagnoses in patients with PPCM that might explain our observations, we queried ICD-9 diagnoses for each individual patient in the PPCM exposure group. Most categories of overlap had fewer than 10 patients; for example, fewer than 10 out of 58 patients with PPCM (6.9%) had overlapping outcome diagnoses (for HF/stroke and HF/MI). No single PPCM exposure had all three outcome diagnoses. Among patients with hypertensive disorder of pregnancy subtypes, GDM, preterm birth, and IUGR exposures, the percentage of patients with more than one outcome diagnosis ranged from 5.74% to 11.37%. Therefore, the majority of observed outcomes were separate cardiovascular events.

We then asked whether heart failure could be a mediator of MI and stroke among patients with PPCM and overlapping outcome diagnoses. We re-fitted our MI and stroke models with heart failure as a time-dependent covariate. With heart failure as a mediator, the risk of PPCM for MI decreased from 13.0 (95% CI 4.1-40.9) to 4.6 (95% CI 1.4-14.8) and the risk of PPCM for stroke decreased from 7.6 (95% CI) to not significant (2.7 95% CI 0.8–8.5). Therefore, although there were novel significant associations between PPCM and MI and PPCM and stroke, heart failure mediated much of those associations.

Finally, we analyzed outcomes by MI and stroke subtypes (**Table 3**). Based on ICD-9 coding, we found that the predominant type of MI experienced within the follow-up period was MICAD, and the most common type of stroke was ischemic stroke.

## Discussion

### Summary of Findings

In this retrospective cohort study of over 1.6 million deliveries representing nearly every completed pregnancy in California over a 5-year period, we report the following: i) PPCM is associated with risks for MI and stroke in addition to (and partly mediated by) HF; ii) HDP subtypes were associated with risks for MI, stroke, and HF; iii) subtypes of HDP representing a longer duration of hypertension and a higher severity (i.e. chronic HTN and chronic HTN with superimposed preeclampsia) had the greatest magnitudes of risks for MI, stroke and HF; iv) GDM, preterm birth, and IUGR were independently associated with risks for future CVD when accounting for PPCM and HDP; and v) PPCM and HDP were independently associated with CVD when accounting for these other CVD-related pregnancy complications. Stratification by age and by selected exposures did not significantly change observed hazard ratios, suggesting these findings are robust to potential residual confounding.

We found that the above cardiovascular pregnancy complications were associated with substantial risks for MI, HF, and stroke even within the first six years post-delivery. In contrast to long-held belief that heart disease largely affects women who are long past their child-bearing years, we demonstrate that the presence of cardiovascular complications of pregnancy can identify a sub-population of younger women who are at high risk for premature cardiovascular disease. Our findings are relevant given the recent call to attention and study concerning the growing epidemic of MI in younger women by scientific experts(7) and support initiatives designed to follow younger at-risk women more closely for CVD prevention(22).

PPCM is hypothesized to have several different causes, including genetic mutations, fetal autoimmunity, vascular dysfunction, in addition to HDP itself(23-25). Furthermore, PPCM presenting during pregnancy may represent a different clinical entity as compared to PPCM presenting postpartum(12) and may have different underlying genetic underpinnings(13). Despite this, our findings did not suggest that intrapartum and postpartum PPCM as coded in HCUP conferred differing risks for CVD. Despite overlapping pathophysiology between PPCM and HDP(26), PPCM has a relatively greater magnitude of risk for incident CVD than any of the HDP subtypes studied.

Given that preterm birth, IUGR, and GDM have demonstrated associations with CVD(27-29) and are commonly seen with HDP and PPCM, we studied these as secondary exposures and did find that each was significantly associated with one or more subtypes of CVD, confirming findings from prior investigations. HDP is a common reason for medically indicated preterm delivery. Several lines of evidence suggest that a common pathology in many cases of preterm birth and IUGR, as well as subtypes of HDP and PPCM and HDP, is vascular dysfunction of varying chronicity and severity.(30-34) GDM is a risk factor for future diabetes and its attendant microvascular and macrovascular effects(35) which may explain the associations seen between GDM and stroke, as well as HF.

### Strengths and Limitations

As a statewide record, California HCUP provided generalizable data from a large and ethnically diverse population and has been used to uncover several insights into cardiovascular disease(18,36,37). Age, race, and ethnicity in the study cohort were similar to those of national CDC birth records for the same time period(19). HCUP captures over 95 percent of California’s population, which comprises over 38 million people. Large datasets like HCUP can facilitate defining phenotypes at greater resolution, such as the pregnancy complication subtypes analyzed in this paper and in other work from our group(18). In addition to several exposures, we were also able to adjust for several potential confounders, which clarified the effects of each exposure on future cardiovascular risk. Increasing phenotypic resolution at scale can help uncover new associations and inspire new hypotheses on the mechanisms of disease. Here we demonstrate for the first time that PPCM, a relatively rare event, carried risks not only for future HF but also for MI and stroke as well, and that these associations, while mediated in part by HF, are not simply explained by overlap of outcome diagnoses.

In terms of magnitude, PPCM conferred the highest magnitude of hazard ratios for future CVD, followed by chronic HTN with superimposed preeclampsia, and then chronic HTN alone. Whereas PPCM and HDP have been demonstrated to exist on a common pathophysiologic spectrum(11,26), we demonstrate that their respective association with later cardiovascular disease differ in magnitude of risk. Further basic and translational studies are needed to determine the specific pathophysiologic mechanisms that connect PPCM and HDP with later CVD.

Relatively short follow-up time (maximum of 6.84 years) likely resulted in an underestimation of risk along the lifetimes of the individuals studied. Also, with small numbers in some groups after adjustment for several covariates, 95% confidence intervals for several exposure subtypes were wide. Given the significant associations found despite these caveats, the hazard ratios measured in this study may in fact be underestimates of true cardiovascular risk following a pregnancy complicated by PPCM and/or HDP.

Despite using a large dataset like HCUP, our analysis of exposure and outcome subtypes were potentially limited due to misclassification bias and to a lack availability of certain diagnostic codes. We attempted to assess the extent of misclassification bias in our study. HCUP data is anonymized, precluding the ability to perform manual chart review of the ICD-9 coding used to define exposures, covariates, and outcomes. We found that prevalence of preterm birth was similar to national estimates; prevalence of PPCM, and GDM were similar to previous reports in smaller California cohorts in which diagnoses were confirmed by manual chart review(19,21). Literature review of ICD-9 coding for preeclampsia, gestational HTN, and hypertensive disorders of pregnancy shows low sensitivity and high specificity(38) while ICD-9 coding for outcomes like MI, HF, and stroke shows high sensitivity and specificity(39-41). If these trends are also true within HCUP, it would suggest that the hazard ratios reported in this study are likely accurate, but could be underestimates of true risk. Despite these shortcomings, our analysis shows several robust signals indicating increased CVD risk among exposures of interest.

Finally, the lack of certain diagnostic codes limited our ability to perform detailed analysis of outcome subtypes (e.g., HF subtypes) or distinguish between HF class. When defining MI and stroke by outcome subtype, there were not enough numbers to perform statistical analysis; the national HCUP database may provide adequate numbers for such an analysis.

In this study, we demonstrated that PPCM is associated with near-term HF, MI and stroke in women in California, independent of HDP, GDM, preterm delivery and IUGR. Among HDP subtypes, chronicity and severity of HTN increased risk of subsequent HF, MI and stroke. While these results are independent of age, the increased prevalence of exposures and outcomes with age in the face of national birth patterns with women giving birth at older ages mean that we can expect even more PPCM, HDP, and CVD in pregnant women. The importance of pregnancy as a cardiovascular ‘stress test’ will, therefore, only further expand in the future.

## Conclusions

Our findings support close monitoring of women with cardiovascular pregnancy complications for prevention of subsequent, near-term CVD events and begs further study of potential mechanisms underlying the development of these early CVD events.

